# Cross-species Y chromosome function between malaria vectors of the *Anopheles gambiae* species complex

**DOI:** 10.1101/151894

**Authors:** Federica Bernardini, Roberto Galizi, Mariana Wunderlich, Chrysanthi Taxiarchi, Nace Kranjc, Kyros Kyrou, Andrew Hammond, Tony Nolan, Mara N. K. Lawniczak, Philippos Aris Papathanos, Andrea Crisanti, Nikolai Windbichler

**Affiliations:** Department of Life Sciences, Imperial College London, South Kensington Campus, London SW7 2AZ, UK.; Malaria Programme, Wellcome Trust Sanger Institute, Hinxton, Cambridge CB10 1SA, UK.; Section of Genomics and Genetics, Department of Experimental Medicine, University of Perugia, 06132 Perugia, Italy.

**Author notes:** Corresponding authors: Nikolai Windbichler, Address: Department of Life Sciences, Imperial College London, South Kensington Campus, London SW7 2AZ, UK, E-mail address Address: Department of Life Sciences, Imperial College London, South Kensington Campus, London SW7 2AZ, UK.

**Keywords:** Vector genetics, hybrid incompatibility, gene flow, Y chromosome, malaria

## Abstract

Y chromosome function, structure and evolution is poorly understood in many species including the *Anopheles* genus of mosquitoes, an emerging model system for studying speciation that also represents the major vectors of malaria. While the Anopheline Y had previously been implicated in male mating behavior, recent data from the *Anopheles gambiae* complex suggests that, apart from the putative primary sex-determiner, no other genes are conserved on the Y. Studying the functional basis of the evolutionary divergence of the Y chromosome in the gambiae complex is complicated by complete F1 male hybrid sterility. Here we used an F1xF0 crossing scheme to overcome a severe bottleneck of male hybrid incompatibilities and enabled us to experimentally purify a genetically labelled *A. gambiae* Y chromosome in an *A. arabiensis* background. Whole genome sequencing confirmed that the *A. gambiae* Y retained its original sequence content in the *A. arabiensis* genomic background. In contrast to comparable experiments in *Drosophila*, we find that the presence of a heterospecific Y chromosome has no significant effect on the expression of *A. arabiensis* genes and transcriptional differences can be explained almost exclusively as a direct consequence of transcripts arising from sequence elements present on the *A. gambiae* Y chromosome itself. We find that Y hybrids show no obvious fertility defects and no substantial reduction in male competitiveness. Our results demonstrate that, despite their radically different structure, Y chromosomes of these two species of the gambiae complex that diverged an estimated 1.85Myr ago function interchangeably, thus indicating that the Y chromosome does not harbor loci contributing to hybrid incompatibility. Therefore, Y chromosome gene flow between members of the gambiae complex is possible even at their current level of divergence. Importantly, this also suggests that malaria control interventions based on sex-distorting Y drive would be transferable, whether intentionally or contingent, between the major malaria vector species.

## Introduction

Sex chromosomes often play an important role in speciation, though the molecular factors that influence this process remains an area of active investigation (Ellegren 2011). Y chromosome sequence content in heterogametic animals is transmitted in a clonal manner due to the lack of crossing over with the X across some or all of its length. This absence of recombination promotes a progressive genetic degeneration including the accumulation and rapid turnover of repetitive sequences (Charlesworth 1991; Rice 1996; Charlesworth & Charlesworth 2000; Bachtrog 2013). It is generally thought that remaining Y-linked genes represent the remnants of an inexorable process of inactivation, degradation, and gene loss, and only genes with a selectable function, such as the male-determining factor are likely to survive on the Y chromosome. However, while this view appears to hold true for mammals, the birth of new genes on the Y has been described in *Drosophila* and may even dominate the evolution of it’s Y chromosome (Carvalho et al. 2015; Vibranovski et al. 2008). This suggests an evolutionary dynamic that is particular to the Y of different phylogenetic groups. Sex chromosomes play an important role in reproductive isolation, however it is mostly the gene-rich X chromosome that has been implicated with three interrelated patterns of hybrid incompatibility: Haldane’s rule, the large X effect, and the asymmetry of hybrid viability and fertility in reciprocal crosses (Wu et al. 1996; Masly & Presgraves 2007). In a few cases a link between the Y and hybrid incompatibilities has been demonstrated. For example it has been shown that the Y chromosome contributes to reproductive barriers between rabbit subspecies (Geraldes et al. 2008). This can be explained by interactions, involved in determining male dimorphism or fertility, between genes which diverged from allele pairs on the Y and X and which break down in the heterospecific context. Such X-Y chromosome incompatibilities have been demonstrated to contribute to hybrid male sterility in house mice (Campbell & Nachman 2014). Additionally the introgression of heterospecific Y chromosomes in *Drosophila* was found to affect male fertility and alters the expression of 2-3% of all genes in hybrids (Sackton et al. 2011).

The varied picture of the Y’s biological role emerging form work in mammals and *Drosophila* suggests the need for additional studies using other model systems. The *Anopheles* genus, which contains all human malaria-transmitting mosquito species has in recent years received much attention, not only due to its stark medical importance but also as a model system for studying speciation and chromosome evolution. In particular the *Anopheles gambiae* species complex of eight sibling species including the most widespread and potent vectors of malaria in Sub-Saharan Africa offers an excellent platform to further our understanding of the biology of the Y and its possible role in reproductive isolation. To this end, we decided to focus on the two species with prime medical importance, *A. gambiae* and *A. arabiensis,* as they are the most anthropophilic members of the complex with the widest distributions.

*A. gambiae* predominates in zones of forest and humid savannah whereas *A. arabiensis* prevails in arid savannahs and steppes, including those of the South-Western part of the Arabian Peninsula. In the sympatric areas, changes in seasonal prevalence are observed showing an increase in the relative frequency of *A. arabiensis* during the dry season. In areas where the distribution of *A. gambiae* and *A. arabiensis* overlaps, hybrids are detected at extremely low frequency (0.02-0.76%) (Toure et al. 1998; Temu et al. 1997; Mawejje et al. 2013). However, a recent study conducted in Eastern Uganda to investigate hybridization between these species showed that 5% of the samples analysed were hybrid generations beyond F1 (Weetman et al. 2014). F1 male sterility and other post-zygotic isolation mechanisms have been studied in *A. gambiae* and *A. arabiensis* hybrids. In addition to mapping multiple loci contributing to male sterility, Slotman et al. demonstrated, by observing the absence of particular genotypes in backcross experiments, that inviability is caused by recessive factors on the X chromosome of *A. gambiae* incompatible with at least one factor on each autosome in *A. arabiensis* (Slotman et al. 2004).

In the present study we wanted to establish whether the introgression of the *A. gambiae* Y chromosome into an *A. arabiensis* genetic background is possible when selected for in a controlled laboratory setting, whether the Y contributes to reproductive isolation and whether a heterospecific Y, as it has been reported in *Drosophila*, would markedly modulate gene expression patterns, fertility or behavior of Y hybrid males. In addition to basic biological insights our attempt to better understand the biology of the mosquito Y is key for both the development of male-specific traits for genetic control as well as predicting the behavior of such traits in the field.

## Materials & Methods

### Mosquito Strains and Rearing

Wild-type *A. gambiae* and *A. arabiensis* mosquitoes of strains G3 and Dongola were used respectively. Strain G3 was originally colonized from West Africa (MacCarthy Island, The Gambia, in 1975) and obtained from the MR4 (MRA-112). It is considered a hybrid stock with mixed features derived from both *Anopheles gambiae s.s*. and *Anopheles coluzzii*. The Dongola strain of *A. arabiensis* was obtained from the MR4 (MRA-856) and was originally isolated from Sudan. For the introgression experiments we used two independently generated Y-linked *A. gambiae* transgenic strains GY1 (Refered to as YattP in Bernardini et al. 2014) carrying a Pax-RFP marker and GY2 (unpublished Roberto Galizi). Strain G^Y2^ contains a Y linked insertion of construct pBac[3xP3-DsRed]β2-eGFP::I-PpoI-124L (Galizi et al. 2014) also carrying a Pax-RFP marker (no expression from the inactive β2-eGFP::I-PpoI-124L locus is detectable in this strain). Inverse PCR suggests position 17757 on the Y_unplaced collection as a likely insertion site of this construct, however no assembly of the repetitive *A. gambiae* Y chromosome exists. All mosquitoes were reared under standard condition at 28 °C and 80% relative humidity with access to fish food as larvae and 5% (wt/vol) glucose solution as adults. For eggs production, young adult mosquitoes (2–4 days after emergence) were allowed to mate for at least 6 days and then fed on mice. Two days later, an egg bowl containing rearing water (dH2O supplemented with 0.1% pure salt) was placed in the cage. One to two days after hatching, larvae were placed into rearing water containing trays. The protocols and procedures used in this study were approved by the Animal Ethics Committee of Imperial College in compliance with UK Home Office regulations.

### Genetic crosses and fertility assays

Crosses were set up in BugDorm-1 cages with size 30x30x30 cm. Generally 100 female and 100 male mosquitoes were crossed during the introgression experiment, although the number of males varied after the F3 bottleneck and was dependent on the number of male progeny that could be recovered from the previous generation. To assay fertility of the Y-introgressed males, after 11 generation of backcrossing in cage, single crosses in cups were set up. Y-introgressed males were singularly introduced into a cup together with one *A. arabiensis* female. Parallel the same number of cups were set up for *A. arabiensis* males and females as a control. After 6 days of mating and blood feeding of females, eggs were collected from every cup and the hatching rate (number of larvae/number of eggs) relative to every cross was calculated.

### DNA sequencing

Samples for DNA sequencing were 10 adult male A^Y2^ and 2 wild-type control *A. arabiensis* males and were used individually for whole genome sequencing. AY2 males had been mated to *A. arabiensis* females prior to DNA extraction allowing us to establish fertility for 7 of the 10 A^Y2^ males. The DNA libraries were prepared in accordance with the Illumina Nextera DNA guide for Illumina Paired-End Indexed Sequencing. AMPure XP beads were used to purify the library DNA and for size selection after which the resulting libraries were validated using the Agilent 2100 bioanalyzer and quantified using a Qubit 2.0 Fluorometer. Sequencing runs were performed on 6 lanes (2 samples per lane) of an Illumina flowcell (v3) on the HiSeq1500 Illumina platform, using a 2×100 bp PE HiSeq Reagent Kit according to the manufacturer’s recommendations. Raw reads were processed using FastQC (Andrews 2010, available online at: http://www.bioinformatics.babraham.ac.uk/projects/fastqc) and trimmomatic (Bolger et al. 2014).

### Variant calling and read coverage analysis

Reads were aligned to the *A. gambiae* PEST reference genome assembly (AgamP4) using BWA mem (Li 2013, bio-bwa v0.7.5a) and sorted using Samtools (v1.2). We used the MarkDuplicates module from Picard tools (v1.9) to remove PCR duplicates and the genome analysis tool kit (GATK, v3.3) to realign reads around indels (McKenna et al. 2010). First we used the GATK modules HaplotypeCaller and UnifiedGenotyper to call raw SNPs and merged them across the 10 A^Y2^ and 2 A male samples using GenotypeGVCFs. Biallelic SNPs were selected using GATK VariantFiltration and SelectVariants modules following the GATK best practices guideline. For the coverage analysis we used bowtie (v1.1.1), exclusively reporting alignments for reads having only a single reportable alignment and displaying no mismatches. We used the bedtools (v2.25) makewindows tool to generate a sliding window bed file with 5kb windows overlapping by 2.5kb. We then used Samtools bedcov to generate per-window read counts and calculate the group means for the 7 fertile A^Y2^ and 2 A male control samples.

### Analysis of Y signature elements

Paired whole-genome sequencing (WGS) reads were mapped using Bowtie2 (Langmead & Salzberg 2012) with standard parameters (bowtie2 -x [consensus_build] -a -1 [ x_R1] -2 [x_R2] -S [x.sam]) against a collection of consensus sequences of all known Y chromosome loci of *A. gambiae* (Hall et al. 2016). Read counts at every locus were generated with Samtools (Li et al. 2009) and normalized by library size and locus length in FPKM. WGS data in the form of paired Illumina reads corresponding to the control samples from *A. gambiae* males (NCBI SRA: SRR534285) and females (NCBI SRA:SRR534286) and *A. arabiensis* females (NCBI SRA: SRR1504792) were taken from the Hall *et al*. study. We also performed a separate analysis to evaluate Y chromosome satellite DNA abundance in the introgressed male samples, in part because satellites Ag53A, B and D due to their short length could not be appropriately assessed using Bowtie2. We used jellyfish (Marçais & Kingsford 2011) to generate unique 25mers (kmers of 25bp long) from each of the 6 known Y satellites consensus sequences (jellyfish count -C -m 25 [stDNA-locus-x.fasta] -o [output] -c 5 -s 1000000000 -t [cores]). Using the same approach we then generated and counted unique 25mers from each of the aforementioned WGS samples (jellyfish count -C -L 5 -m 25 [x.fastq] -o [output] -c 5 -s 1000000000 -t [cores]) and assessed the abundance of each of the stDNA specific kmers within the sample-specific kmers list, resulting in a table providing abundance of each kmer in each sample (grouped by stDNA locus). Raw kmer counts were normalized by library size (sequencing depth) and for the WGS generated in this study (control and introgressed males) we calculated the median abundance for each kmer in these two groups.

### RNA and small RNA sequencing experiments

Sample for RNA sequencing were generated using the following experimental design. Four cages were set up with 40 *A. arabiensis* wild-type males and 40 wild-type females and four cages with 40 A^Y2^ males and their non-transgenic sibling females. After mating and blood feeding, progeny was collected from the cages. 80 freshly hatched larvae were collected from each of both sets of cages and combined in a single tray for rearing. At the pupal stage males were sexed and screened for the fluorescent marker linked to the Y chromosome. From every tray 18 RFP positive and 18 RFP negative males were collected and placed in separate cages to allow emergence. Three days after emergence males were dissected in order to separate three different tissues the head, the abdominal segments harboring the reproductive tissues and the remainder of the carcass. This experiment was performed in twice resulting in a total number of 8 replicates for both controls and experimental samples. Libraries for total RNA sequencing were prepared using the TruSeq RNA sample preparation kit by Illumina and sequenced on 3 lanes of an Illumina HiSeq 2500 using 2x100 paired end reads. The samples described above were also used for the construction of libraries for small RNA sequencing. Libraries were prepared using the NEBNext Multiplex Small RNA kit for Illumina and sequenced on 3 lanes of a MiSeq using the 1x42 single-read mode.

### Differential expression analysis

Reads were aligned to the *Anopheles arabiensis* genome supplemented with the 25 contigs corresponding to known *A. gambiae* Y loci (Hall et al. 2016) using HISAT2 (Pertea et al. 2016). We then used Stringtie (Pertea et al. 2016) in conjunction with the *Anopheles arabiensis* reference transcriptome version AaraD1.3 to predict novel genes and novel isoforms of known genes which were merged across samples into a combined geneset using Cuffmerge (Ghosh & Chan 2016). Expression of all transcripts was then quantified using Stringtie -B. We used the Ballgown (Pertea et al. 2016) suite to determine transcripts showing significantly different expression levels using a cutoff of p-value <0.05 adjusted for multiple testing, a mean expression greater than 1 FPKM across the samples in the tissue analyzed and a log2 fold-change greater than 1 between the introgressed and control groups. To assign a repetitiveness score the sequence of each transcript was blasted against the *Anopheles gambiae* PEST RepeatMasker library provided by Vectorbase.org. For the small RNA dataset we generated a count matrix using Seqbuster and SeqCluster suites (Pantano et al. 2011) and used DESeq2 (Love et al. 2014) for differential expression analysis and log2 transformation of the count data. Only putative small RNA loci with a mean expression of more than 5 counts across samples were taken into account for this analysis.

### Competition experiments

Matings were performed in BugDorm-1 cages. Females and competing males were allocated in cage with a 1:1:1 ratio. Every experiment was run in triplicate. Crosses were set up as follows: 50 A females x 50 A males + 50 G^Y2^ males, 50 G females x 50 G males + 50 A^Y2^ males, 50 A females x 50 A males + 50 A^Y2^ males, 50 G females x 50 A males + 50 A^Y2^ males. After six days mating and blood feeding females were collected from each experimental replicate and allowed to lay singularly in cups. The number of eggs and hatched larvae was calculated for every cup in order to estimate the hatching rate values. Progeny was screened for 3xP3 RFP to assess paternity and transgene ratio was calculated in order to identify any occurring secondary mating.

## Results

### Experimental introgression of the *A. gambiae* Y chromosome into *A. arabiensis*

We have previously established a number of transgenic strains in which different fluorescent transgenes were inserted onto the *A. gambiae* Y chromosome (Bernardini et al. 2014) (Roberto Galizi unpublished). Limited recombination between the *A. gambiae* sex chromosomes has been suggested to occur in specific genetic backgrounds (Wilkins et al. 2007) but is generally believed to occur at very low frequencies (Mitchell & Seawright 1989; Bernardini et al. 2014; Hall et al. 2016). We concluded, that for our purpose a Y-linked fluorescent transgene would be a reliable tool to track the Y chromosome throughout a multi-generational introgression experiment. It has previously been shown that F1 male hybrids between *A. gambiae* and *A. arabiensis* suffer from complete male sterility, which precludes Y chromosome introgression experiments in a straightforward manner. Our pilot experiments confirmed this finding (Supplementary Table 1A). We employed a F1xF0 crossing strategy (we define this as a scheme where first generation female hybrids are crossed to pure species males) in which transgenic Y *gambiae* males were backcrossed to F1 hybrid females that were in-turn generated using either wild type *gambiae* or *arabiensis* mothers (Figure 1A). We used two different transgenic Y strains herein referred to as G^Y1^ and G^Y2^ that both express an RFP reporter gene driven by the neuronal 3xP3 promoter from different insertion sites on the Y chromosome. In the F2 cross, the hybrid males containing the labeled Y are crossed again to wild type *arabiensis* females, and then third generation hybrid males are crossed to both hybrid females and wild type *arabiensis* females in the attempt to recover offspring. Using this crossing scheme, we encountered a severe bottleneck at generation F3 when males are predicted to have inherited a predominantly *A. gambiae* autosomal genome from their fathers in conjunction with pure *A. arabiensis* genome (including the X chromosome) from their mothers (Figure 1B). We recovered no larvae from >13,000 eggs when backcrossing of either G^Y1^ or G^Y2^ males was carried on from F1 hybrid females originated from *A. gambiae* mothers. In the reverse cross, using F1 hybrid females from *A. arabiensis* mothers, we managed to recover 7 larvae (3 males, 4 females) with strain G^Y2^ in a direct cross to wild type *A. arabiensis* females (Figure 1A). These 3 hybrid males obtained were used to progress the introgression and we continuously maintained backcross purification by crossing hybrid males recovered each generation to wild-type *arabiensis* females for a total of 11 generations to establish the introgressed strains A^Y2^ used for all subsequent experiments in this study. The occurrence of fertile phenotypes in these crosses is a rare event. In additional experiments were either G^Y1^ or G^Y2^ males were crossed to F2 or F3 hybrid females (Figure 1A) the majority of the resulting male progeny (15 larvae of which 7 were male and 34 larvae of which 13 were male) failed to develop into fertile adults.

**Figure 1.**
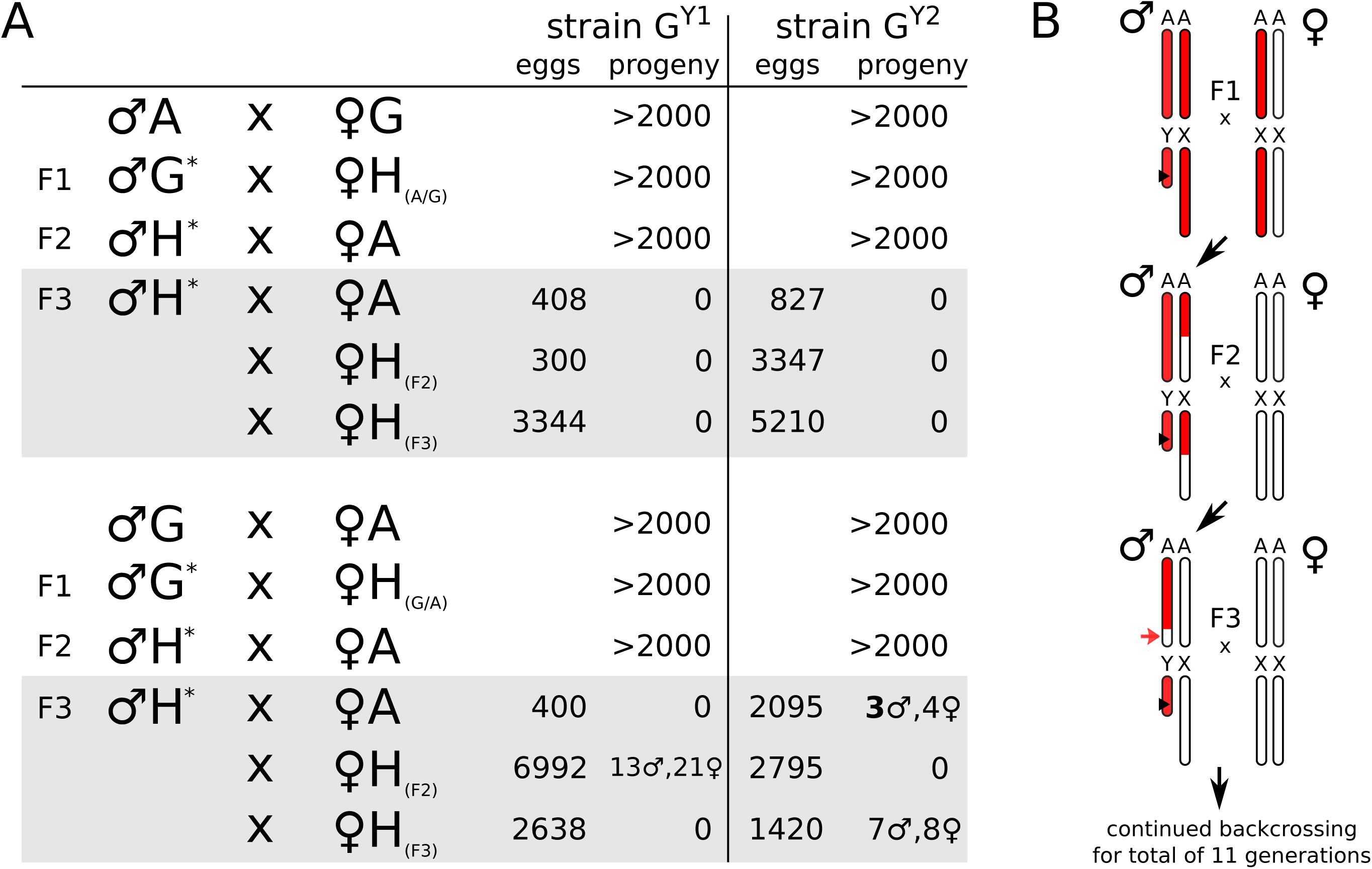
F0xF1 hybrid crossing scheme and resulting progeny. **(A)** Observed number of eggs and progeny arising for each indicated cross where the F1 hybrid females had either an *A. gambiae* mother (top) or an *A. arabiensis* mother (bottom). The asterisk indicates strain G^Y1^ or G^Y2^ respectively. In the bottleneck generation (F3) males were crossed to either pure species arabiensis females (♀A) or crossed to female hybrids with decreasing levels of *A. gambiae* genome content (♀H_(F2)_, ♀H_(F3)_). **(B)** Crossing scheme indicating the *A. gambiae* (red) or *A. arabiensis* (white) genomic contributions in generations F1 to F3 with autosomes represented as a single pair labeled A. In the F1 cross *A. gambiae* males of strains G^Y1^ or G^Y2^ are crossed to hybrid F1 females. Each generation the resulting males are backcrossed to *A. arabiensis* wild-type females. In the bottleneck generation F3 hybrid males harbour an *A. arabiensis* X with a fraction (~25% on average) of autosomal regions expected to be homozygous for the *arabiensis* background (red arrow).

### Fertility of males carrying a heterospecific Y chromosome

We first asked whether A^Y2^ males showed reduced levels of fertility when compared to wild-type *A. arabiensis* males due to the presence of the heterospecific Y chromosome. In order to assay individual males, rather than a population average, we performed single-copula mating experiments of strain A^Y2^ or wild-type males mated to *A. arabiensis* females, measuring the mating rates and oviposition and egg hatching rates of single females. Every generation the males of the largest family were used for establishing the next round of single-copula backcrosses and over the course of 7 generations a total of 344 wild type *A. arabiensis* and 350 A^Y2^ males were assayed. The rationale for this design was to exclude the possibility of *A. gambiae* fertility loci having been retained by selection in introgressed males, because such loci would be expected to segregate in this design. Figure 2 shows a summary of these experiments. Single copula matings are inefficient, in fact less than 25% of females would mate and oviposit under these condition (Figure 2A). No significant difference in the rate of mating was observed between A^Y2^ (22%) and control *A. arabiensis* males (15.3%). Power analysis suggested that the sample size of the successfully mated males would allow for the reliable detection of an effect of medium size (*p*=0.79 for a two-sided t-test, p-value=0.05, d=0.5). We observed that females mated to A^Y2^ males and females mated to *A. arabiensis* wild-type males laid a comparable numbers of eggs (Figure 2B) that had comparable hatching rates (Figure 2C). This analysis indicates that, under laboratory conditions and in the absence of mate choice and male-male competition and taking into account above considerations on power, A^Y2^ males show no significant difference in fertility when compared to wild-type *A. arabiensis* males that retain their native Y chromosome.

**Figure 2.**
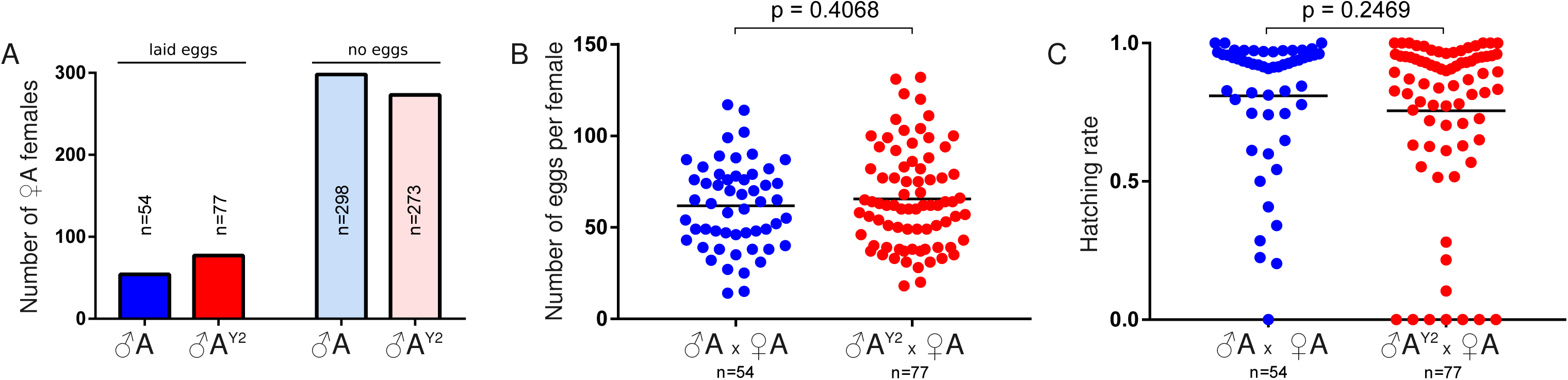
Single copula mating experiments. **(A)** Number of females ovipositing following single matings with A or A^Y2^ males. Counts of eggs **(B)** and hatching larvae **(C)** for each individual family.

As an additional control, we back-crossed A^Y2^ males to *A. gambiae* females. Despite the presence of the *A. gambiae* Y chromosome in these males we expected this experiment to recreate hybrid incompatibility in the form of male infertility in the resulting progeny. Indeed we found hybrid males to be fully sterile (Supplementary Table 1B) thus confirming that X-A incompatibilities are sufficient to explain this phenotype (Slotman et al. 2004).

### Genomic analysis of males with a heterospecific Y chromosome

After n=11 generations of backcrossing, assuming no selection for sections of the *A. gambiae* genome, the expected autosomal genome proportion of the *A. gambiae* donor would be 1/2^n^ or less than 0.05%. Given that *gambiae* genomic regions contributing to hybrid incompatibilities would be selected against in males this is likely an underestimation due to such detrimental haplotypes being selectively removed. To confirm that our backcrossing scheme had eliminated the *A. gambiae* autosomal genome but retained the *A. gambiae* Y chromosome we performed whole genome DNA sequencing from 10 A^Y2^ males and 2 wild-type control males of our *A. arabiensis* lab colony. To determine whether introgressed males were fertile, they were singularly mated to wild-type arabiensis females before their genomic DNA was extracted. WGS reads were mapped to the *A. gambiae* genome to identify genomic regions with fixed allele differences between introgressed and control groups by confining our analysis to biallelic SNPs. No assembly of the *A. gambiae* Y exists, however the PEST assembly includes the Y_unplaced sequence collection that includes ~230kB of unscaffolded contigs that have been assigned to the Y. Our analysis (Table 1) showed that the autosomes of introgressed-Y and pure species contained a small number of differentially represented SNPs comparable in number to the X chromosome which, since it is replaced in every backcross generation, serves as a background control. In contrast, the majority of differentially represented SNPs (74.7% of the total number of differential fixed SNP and 35.6% of the total number of SNPs on the Y) arose from reads mapping to the Y_unplaced portion of the *A. gambiae* genome that represents less than 0.1% of the total genome assembly. Within the Y_unplaced collection we found that most SNPs mapped to the largest contig, which also contains the male-determine gene (Supplementary Figure 1). In addition 12.7% of of the total number of SNPs arose from the UNKN collection (unassigned contigs) that is also expected to contain a number of unassigned Y sequences and repetitive elements. We performed an additional sliding-window analysis where we considered only reads mapping uniquely to the *A. gambiae* genome and allowed for no mismatches. The rationale was that, given the observed levels of divergence between these genomes, perfectly matching reads are expected to predominantly map to the genome of origin. When comparing the mean number of reads of A^Y2^ fertile introgressed males and the samples of the *A. arabiensis* control group we find that the Y_unplaced collection experiences significant coverage only in A^Y2^ males as do parts of the UNKN collection. For the autosomes and the X, few windows accrue a significant number of reads and we find no substantial differences between the groups in the direction of A^Y2^, with the possible exception of an intergenic region on chromosome 2L (Supplementary Figure 2). This suggests that, apart from the Y, both groups have a similar *A. arabiensis* background and we concluded that the transgenic *A. gambiae* Y had been successfully purified in an *A. arabiensis* genomic background, although our data can’t rule out that some fraction of *A. gambiae* genomic DNA other than the Y chromosome persists in the A^Y2^ strain.

**Table 1.** Analysis of single nucleotide polymorphisms between A and A^Y2^ males

| chromosome | base pairs | total number<br>of variants | biallelic SNPs w/o<br>missing data | differentially fixed SNPs |  |
| --- | --- | --- | --- | --- | --- |
|  |  |  |  | number | % |
| X | 24393108 | 1040639 | 583061 | 27 | 0.0046 |
| 2L | 49364325 | 2180463 | 1464616 | 42 | 0.0029 |
| 2R | 61545105 | 2233941 | 1573367 | 12 | 0.0008 |
| 3L | 41963435 | 1702574 | 1159627 | 27 | 0.0023 |
| 3R | 53200684 | 2480254 | 1653367 | 6 | 0.0004 |
| Mt | 15363 | 23 | 21 | 0 | 0 |
| Unknown | 42389979 | 749098 | 351870 | 115 | 0.0327 |
| Y_unplaced | 237045 | 4180 | 1902 | 677 | 35.5941 |

### Analysis of Y chromosome sequence content in introgressed males

Y chromosomes of *A. gambiae* and *A. arabiensis* have been shown to differ dramatically in their structure and sequence content (Hall et al. 2016). To assess whether introgression of the *A. gambiae* Y chromosome into the *A. arabiensis* genome coincided with any detectable structural re-arrangements of the Y, for example the selective elimination of sequences that would be detrimental in hybrids, we mapped DNA reads from all A^Y2^ individuals as well as pooled control datasets from Hall et al. against a collection of known Y chromosome genes and repeats of *A. gambiae*. This collection includes consensus sequences of all known and putative genes, repetitive elements as well as satellite DNA (Hall et al. 2016). We assessed normalized read depth at each locus (Fragments Per Kilobase of transcript per Million mapped reads) as a proxy for copy number of each sequence element in a given background. This analysis is complicated by the occurrence of autosomal copies of many of these elements as well as possible variation between Y chromosome isolates. Figure 3 shows normalized read counts for these elements in strain A^Y2^ plotted versus males and females of both *A. gambiae* and *A. arabiensis*. We observe an excellent correlation in the representation of these Y signature sequence elements between of *A. gambiae* males and A^Y2^ males. Because these signature elements are derived from and to some degree specific to the *A. gambiae* Y chromosome, they were under-represented in *A. arabiensis* control males and in females of both species. Interestingly, the AgY280 satellite shows an approximately 15x higher coverage in *A. gambiae* males compared to the A^Y2^ strain. However this satellite is known to be highly variable between Y isolates (Hall et al. 2016). In order to better assess the content of DNA satellites, in particular satellites Ag53A, B and C that are too short for standard read mapping, we performed an additional analysis based on kmer counts in the read libraries. This analysis showed that A^Y2^ males resemble *A. gambiae* males with regards to the content of DNA satellites attributed to the Y (Supplementary Figure 3). Together these analyses therefore confirm the presence of the *A. gambiae* Y chromosome in the introgressed strain and suggest that the introgression experiment did not impact in any evident way the original content of this Y chromosome.

**Figure 3.**
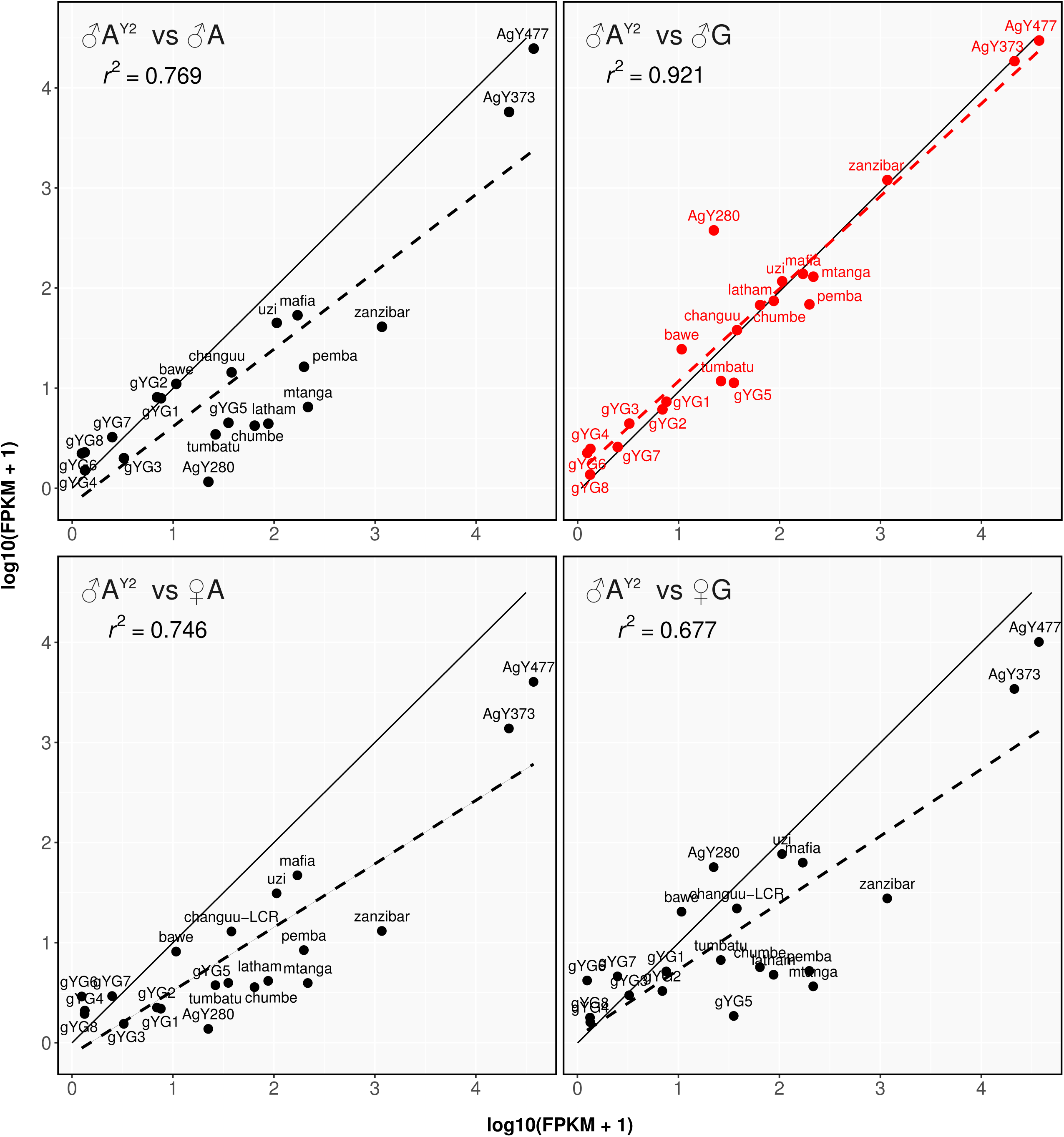
Analysis of the content of the introgressed Y chromosome. The plots show the number of normalized reads mapping to the *A. gambiae* Y chromosome reference loci calculated as log_10_ transformed FPKM values for A^Y2^ males on the x-axis compared to either wild type *A. arabiensis* males (top left panel), wild-type *A. gambiae males* (top right panel), wild-type *A. arabiensis* females (bottom left panel) and *A. gambiae* females (bottom right panel) on the y-axis. The dashed linear regression line and associated r^2^ coefficient indicate the best correlation in read counts of signature Y elements between A^Y2^ and *A. gambiae males*.

### Transcriptomic analysis of males carrying a heterospecific Y chromosome

We next asked to what extent the presence of a heterospecific Y chromosome would alter the expression of autosomal or X-linked genes of the *A. arabiensis* background. This could occur by (i) the action of *gambiae* Y trans-acting factors or absent *A. arabiensis* Y trans-acting factors, (ii) by Y sequences that recruit cellular trans-acting factors and/or modulate the chromatin structure and epigenetic state of other chromosomes or (iii) by the activation of mobile genetic elements present on the *A. gambiae* Y that trigger a cellular response in X or autosomal sequences. We focused our analysis on 3 tissues, the head, the terminal abdominal segments containing the reproductive tissues and the remainder of the carcass. We performed RNA sequencing of a total of 8 control and 8 experimental samples for each tissue from A^Y2^ and wild-type males that had been reared in the same larval tray and that were sexed and separated at the pupal stage. Paired-end reads were mapped against the 1214 *A. arabiensis* genomic scaffolds supplemented by 25 consensus sequences corresponding to known *A. gambiae* Y loci previously mentioned, as well as the reporter gene construct. Due to the incomplete annotation of the *A. arabiensis* genome we performed an isoform level analysis were we first predicted novel genes and novel isoforms of known genes across all samples. Gene-level expression of the experimental groups as well as sample relationships are summarized in Supplementary Figure 4 and Supplementary File 1. Finally, in order to indicate whether differentially expressed transcripts potentially represented known mobile elements or repetitive DNA arising from the Y (but matching paralogous sequences present on the *A. arabiensis* scaffolds which could thus be misreported as differentially expressed) we blasted each predicted transcript to the *A. gambiae* repeat library and assigned a repetitiveness score. We then predicted differential expression of transcripts between A^Y2^ and wild-type males (Figure 4, Supplementary File 2). Few transcripts were expressed significantly lower in A^Y2^ males. This is partially expected because *A. arabiensis* genomic scaffolds are derived from the DNA of females (Neafsey et al. 2015) and it is thus not possible to identify any *A. arabiensis* Y-linked genes that would have fallen into this class. However, it also indicates that few if any endogenous genes are down-regulated as a result of the presence of the heterospecific Y chromosome. In contrast a number of transcripts displayed significantly higher levels of expression in A^Y2^ males. However, the majority of these correspond to the known *A. gambiae* Y loci (purple triangles in Figure 4) and their expression is also summarized in Supplementary figure 5. Of the remaining upregulated *A. arabiensis* transcripts (colored circles in Figure 4) the majority had a high repetitiveness score i.e. significant homology to *A. gambiae* repeats. A manual homology search using all *A. arabiensis* transcript sequences passing the significance threshold and cut-off for expression confirmed that virtually all differentially expressed transcripts are related to repetitive DNA. In addition to the above analysis we also measured small RNA expression in these tissues finding no evidence for the differential expression of small non-coding RNAs between A^Y2^ and wild-type males (Supplementary Figure 6). While it is possible that such effects could occur in other developmental stages or under specific environmental conditions we find little evidence that the heterospecific Y-chromosome markedly affects expression of the non-repetitive, autosomal or X-linked gene repertoire of the *A. arabiensis* genome.

**Figure 4.**
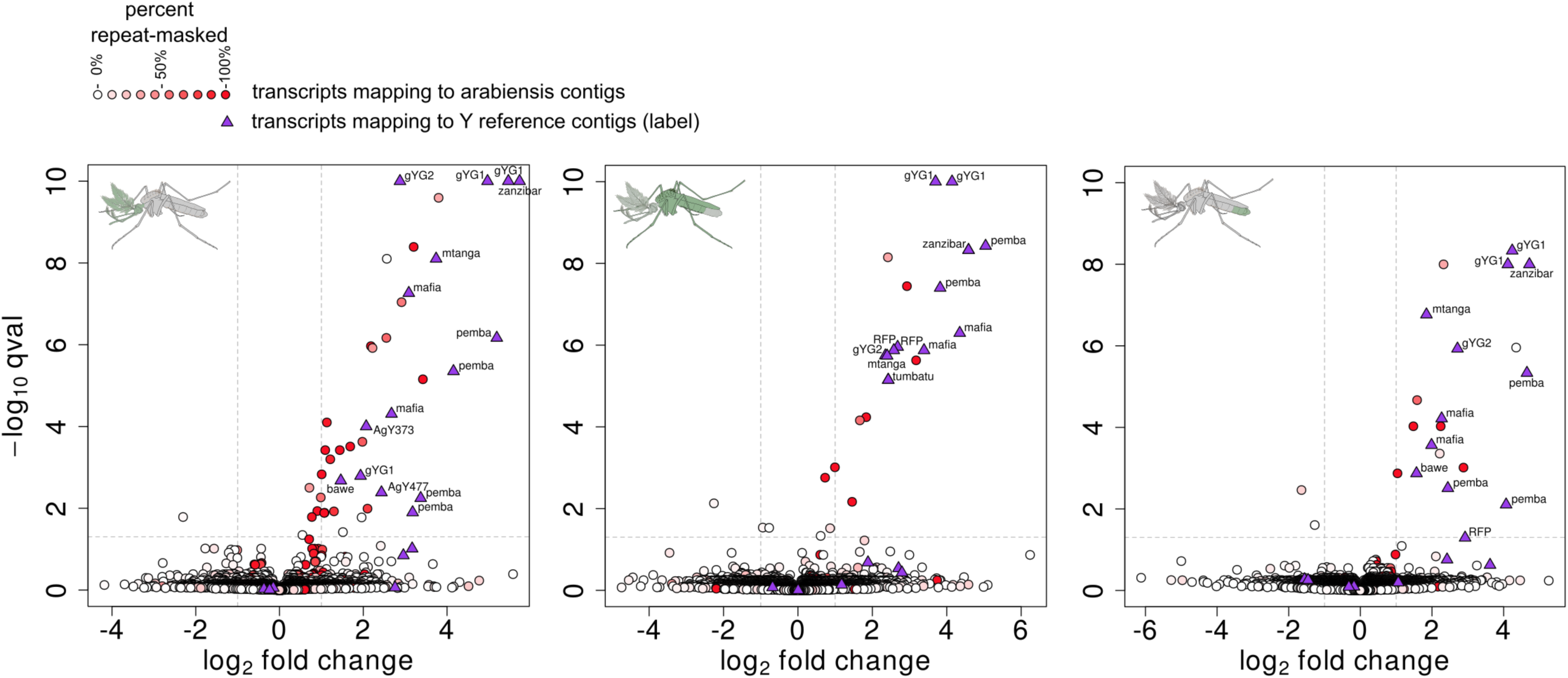
Differential expression analysis by RNA-seq. Volcano plots showing log_2_ fold-change values (x-axis) by -log_10_ corrected p-values (y-axis) for all transcripts between introgressed A^Y2^ males and *A. arabiensis* control males. The analysis was performed separately for the head (left panel), the carcass (middle panel) and the abdominal segments containing the reproductive tract (right panel). Transcripts derived from *A. arabiensis* scaffolds are represented as circles and colored based on the percentage of their sequence masked by sequences in the *A. gambiae* repeats library. Transcripts from the reference set of *A. gambiae* Y loci are indicated by purple triangles and the name of the signature locus. Dashed lines represent the thresholds used for adjusted p-value (p < 0.05) and log_2_ fold change (>1.0).

### Female mate-choice and male competition experiments

Although we find no strong effect of the heterospecific Y chromosome on the transcriptome and fertility of individual A^Y2^ males it is possible that Y-linked sequences do play a role in male fitness or female choice which would also limit the practical use of introgressed Y-linked traits e.g. for vector control. In order to test this hypothesis we set up a panel of competitive mating experiments where two strains of males were allowed to compete for mating with females in population cages (Figure 5). Both wild-type *A. arabiensis* and A^Y2^ males performed substantially worse (winning only 13.7% and 11.1% of matings respectively) than *A. gambiae* males in competition experiments for either *A. gambiae* or *A. arabiensis* females. These findings confirms previous observations (Schneider et al. 2000) and demonstrates that, irrespective of the female type *gambiae* males are superior to *A. arabiensis* males in the lab setting. In contrast A^Y2^ males were only slightly less competitive compared to wild type *arabiensis* males winning ~40% of matings with *arabiensis* females (p=0.0146) and no significant difference was observed when they competed against arabiensis males for *gambiae* females (p=0.156). The particular set up of the experiments also allowed to score for secondary mating measured by the percentage of transgenic male progeny. With the possible exception of one case, re-mating (showing a significant deviation from a 50% transgene ratio in the progeny) was not observed in these experiments. Overall a slight reduction in male competitiveness was observed which could relate, in addition to the effect of the heterospecific Y, to inbreeding or fitness costs associated with expression of the fluorescent marker gene. Importantly, in all cases no significant difference to the wild type *A. arabiensis* males observed in terms of the number of laid and hatching eggs from matings with A^Y2^ males confirming our previous analysis.

**Figure 5.**
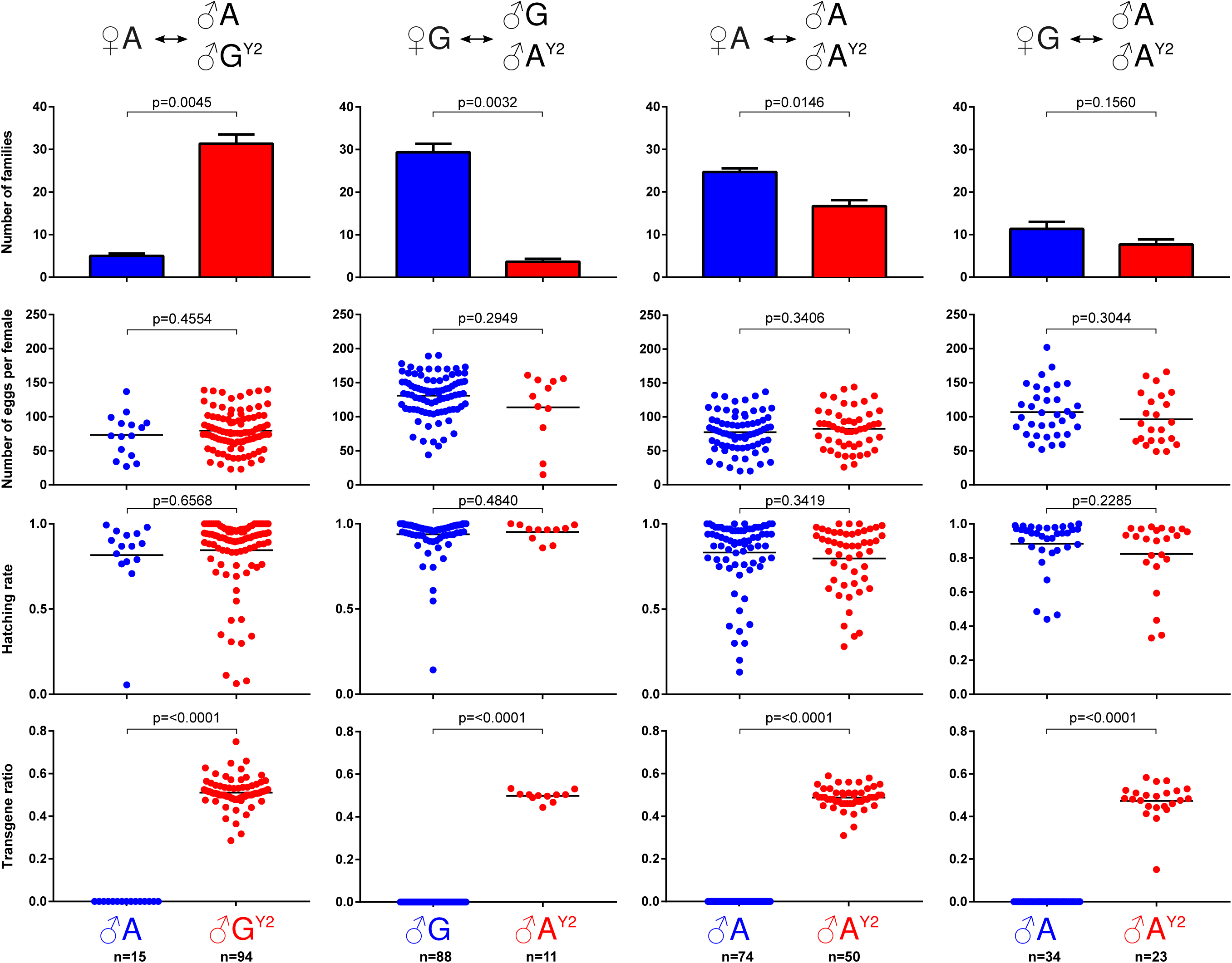
Mating competition experiments. The genotypes of the females and the two types of competing males is indicated at the top for each experiment. The second, third and fourth row of panels show the number of eggs laid by individual females, the hatching rate for each family as well as the ratio of transgenic to wild-type larvae respectively. P-values were calculated using Welch’s two-tailed t-test.

## Discussion

In a classic study Slotman et al. mapped quantitative trait loci related to male sterility in hybrids between *A. gambiae* and *A. arabiensis* and at least five or six sterility factors were detected in each of the two species. The X chromosome was found to have a disproportionately large effect on male hybrid sterility (Slotman et al. 2004), which is likely related to divergent alleles present within multiple fixed chromosomal rearrangements on the X. A possible role of the Y chromosome in hybrid incompatibilities was suggested by the authors but not followed up on experimentally. Using an F1xF0 crossing scheme we generated F3 males with an *arabiensis* X chromosome and a set of *arabiensis* autosomes as well as an *A. gambiae* Y chromosome. The second set of autosomes is expected to contain on average 75% *A. gambiae* sequences. The majority of such F3 males were expected to be sterile; however we hypothesized that it should be possible to select a small fraction of fertile males that lacked the *A. gambiae* incompatibility loci that cause sterility when interacting with the *A. arabiensis* background. Indeed, we recovered 56 larvae out of a total 29,776 eggs laid (0.18%) from pooled backcrosses of ~600 F3 males sampled.

Surprisingly, after multiple generations of backcross purification we found that the *A. gambiae* Y chromosome does not markedly influence male fertility, fitness or gene expression in Y hybrids. This rules out the Y chromosome as a major factor contributing to hybrid incompatibilities and, more importantly, it is in stark contrast to findings in the *Drosophila* model (Sackton et al. 2011). Despite the Y’s relative paucity of genes in the fly even intraspecific Y chromosome variants profoundly affect the expression a substantial number of genes located on the X or the autosomes. The introgression of heterospecific Y chromosomes in *Drosophila*, to the extent that it is even possible (Johnson et al. 1993), has consequently been found to markedly affect male fertility and gene expression in interspecific hybrids. In a *D. simulans* background the *D. sechellia* Y has little effect on viability, but it reduced male fecundity by 63% as well as sperm competitiveness (*D. simulans/sechellia* divergence time has been estimated at only 0.25 Myr). Y introgression differentially affected genes involved in immune function and spermatogenesis suggesting a trade-off in investment between these processes. However, it has also been suggested that a significant part of the observed effect in *Drosophila* may be attributed to the ribosomal DNA (rDNA) clusters that are abundant on both the X and the Y in the fly. Importantly, in the *A. gambiae* strain G3 we used for this study (unlike some strains such as (Wilkins et al. 2007)) and in *A. arabiensis* the rDNA is located exclusively on the X-chromosome and thus does not confound introgression experiments the same way. Our results may thus be more indicative than similar experiments in *Drosophila* of the true effect a gene-poor heterospecific Y exerts on the genome of a related species.

It is possible that more subtle effects of the introgressed Y, not detectable in our experimental setup, exist. Future work could involve hybrid performance testing in mating swarms or under semi-field conditions. The fact however that, despite its radically different structure and an estimated divergence time of ~1.85 Myr or more than 7 times that between *D. simulans* and *D. sechellia*, the *A. gambiae* Y seems to be able to fully replace the *A. arabiensis* Y, suggests that in *Anopheles* the Y either carries no important factors that diverged between these two species or that no such factors are present at all. Although, early work had implicated the Anopheline Y chromosome in mating behavior in a study using *A. labranchiae* and *A. atroparvus* species (Fraccaro et al. 1977) our work is more in line with a recent study that leveraged long single-molecule sequencing to determine the content and structure of the Y chromosome of the primary African malaria mosquito, *A. gambiae* in comparison to its sibling species in the gambiae complex (Hall et al. 2016). This study revealed rapid sequence turnover within the species complex with only a single gene, recently shown to act as a male-determining factor (Krzywinska et al. 2016), being conserved and exclusive to the Y in all species. This gene called YG2 (alternatively referred to as Yob) was not detectable in the genome of the more distant mosquito relative *Anopheles stephensi*, overall suggesting rapid evolution of Y chromosome in this highly dynamic genus of malaria vectors.

The role of gene flow between *A. gambiae* and *A. arabiensis* in leading to adaptive introgression and the implications for vector control has been highlighted by Weetman and colleagues (Weetman et al. 2014). Post-F1 gene flow occurs between *A. gambiae* and *A. arabiensis*, and, especially for traits under strong selection, could readily lead to adaptive introgression of genetic variants relevant for vector control. Introgression of the Y chromosome between species is generally viewed as unlikely and markers found on the Y generally show restricted gene flow relative to other loci. However, this contrasts with the recent hypothesis (Neafsey et al. 2015; Hall et al. 2016) of Y gene flow between *A. gambiae* and *A. arabiensis*. Contrary to the established species branching order, the YG2 gene tree suggested that Y chromosome sequences may have crossed species boundaries between *A. gambiae* and *A. arabiensis* (Hall et al. 2016) and the authors suggested that Y chromosome introgression could have predated the development of male F1 hybrid sterility barriers that exist between this pair of species (Hall et al. 2016). Our study lends support to this hypothesis as it experimentally demonstrates the possibility of a cross-species Y chromosome transfer and shows that the Y is functional in this context.

Our findings thus also suggest that Y-linked genetic traits generated in *A. gambiae* could be transferred to its sister species. For example strain A^Y1^ carries a site-specific docking site that now also allows the generation of male-exclusive genetic traits in *A. arabiensis*. Recent progress towards the elusive goal of efficient sex-ratio distortion by a driving Y chromosome (Galizi et al. 2014; Bernardini et al. 2014) could lead to the development of invasive distorter traits in *A. gambiae* that may then be transferred to sibling species. This could be done deliberately in the lab as we have demonstrated here, but it could also occur contingently in the wild after a large-scale release of transgenic males. While one should always be wary extrapolating lab studies to conditions in the field, the notion that the Y chromosome does not represent a genetic barrier for gene flow between two members of the *A. gambiae* species complex (*A. gambiae* and *A. arabiensis*) should inform the design and implementation of genetic control interventions based on transgenic mosquitoes.

**Supplementary Table 1.** F1 hybrid male crosses and observed progeny. G: *A. gambiae*; A: *A. arabiensis*; H: hybrid; A^Y2^: introgressed strain.

**Supplementary Figure 1.** Allele frequency and position of biallelic SNPs predicted between 2 wild-type *A. arabiensis* and 10 A^Y2^ males within the Y_unplaced sequence collection. Dashed lines indicate breaks between Y_unplaced sequence contigs whose order is not continuous. The black triangle indicates the presumed location of the transgene construct.

**Supplementary Figure 2.** Genome-wide read density maps showing the mean number of reads of the introgressed (7 fertile A^Y2^ males) and *A. arabiensis* control groups (2 A males). The x-axis represents the chromosomal position and the y-axis the normalized number of reads per 5kb sliding window with 2.5kb overlaps for a maximum of 10^6^ reads per window. Only uniquely mapping reads and those lacking mismatches were considered for this analysis. The panels show the five major chromosome arms in the *A. gambiae* genome assembly plus the UNKN and Y_unplaced sequence collections. The black arrow indicates the presumed location of the transgene construct within the Y_unplaced collection.

**Supplementary Figure 3.** Violin plot showing the kmer abundance analysis for 6 *Anopheles gambiae* Y-linked DNA satellites represented as normalized log10 transformed count values in males (blue) and females (red) of control and introgressed strains.

**Supplementary Figure 4.** (A) Heatmaps of all RNAseq sample libraries based on the Euclidean distance of each library summed across all transcripts. (B) Distribution of gene-level FPKM expression values across the experimental groups and the 3 tissues analyzed.

**Supplementary Figure 5.** RNA expression levels from *Anopheles gambiae* Y signature elements as fragments per kilobase of element per million fragments mapped (FPKM) values represented as log_10_ transformed FPKM values for males of the introgressed (A^Y2^) and control groups (A) calculated for each of the analyzed tissues.

**Supplementary Figure 6.** Analysis of small RNAs. The left panels show MA plot representations of the expression of small non-coding RNAs. The analysis was performed separately for the head (A), the carcass (B) and the segments containing the male reproductive tract (C). No small RNA locus passed the significance (adjusted p-value <0.05) and log2 fold change (>1.0) threshold that would indicate differential expression between introgressed and control samples. The right panels indicate the size distribution of small RNAs analyzed on the x-axis and the number of unique RNA species identified of each length is shown on the y-axis. The color of the bars indicates the total number of reads for all the RNA species in each bin as defined in the legends.

## References

Bachtrog, D., 2013. Y chromosome evolution: emerging insights into processes of Y chromosome degeneration. Nature Reviews Genetics, 14(2), pp. 113–124.

Bernardini, F. et al., 2014. Site-specific genetic engineering of the Anopheles gambiae Y chromosome. Proceedings of the National Academy of Sciences, 111(21), p.201404996. Available at: http://www.pnas.org/content/early/2014/05/08/1404996111%5Cnhttp://www.ncbi.nlm.nih.gov/pubmed/24821795%5Cnhttp://www.pnas.org/content/early/2014/05/08/1404996111.abstract.html?etoc%5Cnhttp://www.pnas.org/content/early/2014/05/08/1404996111.full.pdf.

Bolger, A.M., Lohse, M. & Usadel, B., 2014. Trimmomatic: a flexible trimmer for Illumina sequence data. Bioinformatics, 30(15), pp.2114–2120. Available at: http://www.ncbi.nlm.nih.gov/pmc/articles/PMC4103590/.

Campbell, P. & Nachman, M.W., 2014. X-Y interactions underlie sperm head abnormality in hybrid male house mice. Genetics, 196(4), pp. 1231–1240.

Carvalho, A.B. et al., 2015. Birth of a new gene on the Y chromosome of [i]Drosophila melanogaster[/i]. Proceedings of the National Academy of Sciences of the United States of America, 112(40), pp. 12450–12455. Available at: http://www.pnas.org/content/112/40/12450.full.

Charlesworth, B., 1991. Evoluton of Sex Chromosomes. Science, 251, pp.1030–1033.

Charlesworth, B. & Charlesworth, D., 2000. The degeneration of Y chromosomes. Philosophical Transactions of the Royal Society B-Biological Sciences, 355(1403), pp. 1563–1572. Available at: http://eutils.ncbi.nlm.nih.gov/entrez/eutils/elink.fcgi?dbfrom=pubmed&id=11127901&retmode=ref&cmd=prlinks%5Cnpapers3://publication/doi/10.1098/rstb.2000.0717.

Ellegren, H., 2011. Sex-chromosome evolution: recent progress and the influence of male and female heterogamety. Genetics, 12(February), pp. 157–66. Available at: http://www.ncbi.nlm.nih.gov/pubmed/21301475.

Fraccaro, M. et al., 1977. Y chromosome controls mating behaviour on Anopheles mosquitoes. Nature, 265, pp.326–328.

Galizi, R. et al., 2014. A synthetic sex ratio distortion system for the control of the human malaria mosquito. Nature communications, 5, p.3977. Available at: http://www.nature.com/ncomms/2014/140610/ncomms4977/full/ncomms4977.htm.

Geraldes, A. et al., 2008. Reduced introgression of the Y chromosome between subspecies of the European rabbit (Oryctolagus cuniculus) in the Iberian Peninsula. Molecular Ecology, 17(20), pp.4489–4499.

Ghosh, S. & Chan, C.-K.K., 2016. Analysis of RNA-Seq Data Using TopHat and Cufflinks. In D. Edwards, ed. Plant Bioinformatics: Methods and Protocols. New York, NY: Springer New York, pp. 339–361. Available at: http://dx.doi.org/10.1007/978-1-4939-3167-5_18.

Hall, A.B. et al., 2016. Radical remodeling of the Y chromosome in a recent radiation of malaria mosquitoes. Proceedings of the National Academy of Sciences, 113(15), p.201525164. Available at: http://www.pnas.org/content/113/15/E2114.abstract.

Johnson, N.A. et al., 1993. The Effects of Interspecific Y Chromosome Replacements on Hybrid Sterility within the Drosophila Simulans Clade. Genetics, 135(2), pp.443–453. Available at: http://www.ncbi.nlm.nih.gov/pmc/articles/PMC1205647/.

Krzywinska, E. et al., 2016. Research | reports 30., 2927(1979), pp. 1431–1434.

Langmead, B. & Salzberg, S.L., 2012. Fast gapped-read alignment with Bowtie 2. Nat Meth, 9(4), pp.357–359. Available at: http://dx.doi.org/10.1038/nmeth.1923.

Li, H. et al., 2009. The Sequence Alignment/Map format and SAMtools. Bioinformatics, 25(16), pp.2078–2079. Available at: http://www.ncbi.nlm.nih.gov/pmc/articles/PMC2723002/.

Love, M.I., Huber, W. & Anders, S., 2014. Moderated estimation of fold change and dispersion for RNA-seq data with DESeq2. Genome Biology, 15(12), p.550. Available at: http://dx.doi.org/10.1186/s13059-014-0550-8.

Marçais, G. & Kingsford, C., 2011. A fast, lock-free approach for efficient parallel counting of occurrences of k-mers. Bioinformatics, 27(6), pp.764–770. Available at: http://dx.doi.org/10.1093/bioinformatics/btr011.

Masly, J.P. & Presgraves, D.C., 2007. High-resolution genome-wide dissection of the two rules of speciation in Drosophila. PLoS Biology, 5(9), pp. 1890–1898.

Mawejje, H.D. et al., 2013. Insecticide resistance monitoring of field-collected anopheles gambiae s.l. populations from jinja, eastern uganda, identifies high levels of pyrethroid resistance. Medical and Veterinary Entomology, 27(3), pp.276–283.

McKenna, A. et al., 2010. The Genome Analysis Toolkit: A MapReduce framework for analyzing next-generation DNA sequencing data. Genome Research, 20(9), pp. 1297–1303. Available at: http://www.ncbi.nlm.nih.gov/pmc/articles/PMC2928508/.

Mitchell, S.E. Insects A.M. and A.R.L.U.-A.G.F. & Seawright, J.A., 1989. Recombination between the X and Y chromosomes in Anopheles quadrimaculatus species A. The Journal of heredity (USA). Available at: http://agris.fao.org/agrissearch/search.do?recordID=US9022071#.WBNpAXy_oNs.mendeley [Accessed October 28, 2016].

Neafsey, D. et al., 2015. Highly evolvable malaria vectors: the genomes of 16 Anopheles mosquitoes. Science, 347(6217), pp. 1258522–1258522. Available at: http://www.sciencemag.org/cgi/doi/10.1126/science.1258522.

Pantano, L., Estivill, X. & Martí, E., 2011. A non-biased framework for the annotation and classification of the non-miRNA small RNA transcriptome. Bioinformatics, 27(22), pp.3202–3203. Available at: http://bioinformatics.oxfordjournals.org/content/27/22/3202.abstract.

Pertea, M. et al., 2016. Transcript-level expression analysis of RNA-seq experiments with HISAT, StringTie and Ballgown. Nat. Protocols, 11(9), pp. 1650–1667. Available at: http://dx.doi.org/10.1038/nprot.2016.095.

Rice, W.R., 1996. Evolution of the Y sex in animals: Y chromosomes evolve through the degeneration of autosomes. BioScience, 46(5), pp.331–343.

Sackton, T.B. et al., 2011. Interspecific Y chromosome introgressions disrupt testis-specific gene expression and male reproductive phenotypes in Drosophila. Pnas, 108(41), pp. 17046–17051.

Schneider, P., Takken, W. & McCall, P.J., 2000. Interspecific competition between sibling species larvae of Anopheles arabiensis and An. gambiae. Medical and Veterinary Entomology, 14(2), pp. 165–170.

Slotman, M., Della Torre, A. & Powell, J.R., 2004. The genetics of inviability and male sterility in hybrids between Anopheles gambiae and An. arabiensis. Genetics, 167(1), pp.275–287.

Temu, E.A. et al., 1997. Detection of hybrids in natural populations of the Anopheles gambiae complex by the rDNA-based, PCR method. Annals of tropical medicine and parasitology, 91(8), pp.963–965.

Toure, Y.T. et al., 1998. The distribution and inversion polymorphism of chromosomally recognized taxa of the Anopheles gambiae complex in Mali, West Africa. Parassitologia, 40(4), pp.477–511.

Vibranovski, M.D., Koerich, L.B. & Carvalho, A.B., 2008. Two new Y-linked genes in Drosophila melanogaster. Genetics, 179(4), pp.2325–2327.

Weetman, D. et al., 2014. Contemporary gene flow between wild An. gambiae s.s. and An. arabiensis. Parasites & Vectors, 7(1), p.345. Available at: http://parasitesandvectors.biomedcentral.com/articles/10.1186/1756-3305-7-345.

Wilkins, E.E., Howell, P.I. & Benedict, M.Q., 2007. X and Y chromosome inheritance and mixtures of rDNA intergenic spacer regions in Anopheles gambiae. Insect Molecular Biology, 16(6), pp.735–741. Available at: http://dx.doi.org/10.1111/j.1365-2583.2007.00769.x.

Wu, C.I., Johnson, N.A. & Palopoli, M.F., 1996. Haldane’s rule and its legacy: Why are there so many sterile males? Trends in Ecology and Evolution, 11(7), pp.281–284.

